# Childhood location correlates with epigenetic age and methylation stability in British-Bangladeshi migrants

**DOI:** 10.1101/2020.09.19.304808

**Authors:** Reinhard Stöger, Minseung Choi, Gregory Leeman, Richard D. Emes, Khurshida Begum, Philippa Melamed, Gillian R. Bentley

**Affiliations:** School of Biosciences, University of Nottingham, LE12 5RD, United Kingdom; School of Medicine, Stanford University, USA; School of Veterinary Medicine and Science, University of Nottingham, LE12 5RD, United Kingdom; Advanced Data Analysis Centre, University of Nottingham, LE12 5RD, United Kingdom; Department of Anthropology, Durham University, Durham, United Kingdom; Faculty of Biology, Technion-Israel Institute of Technology, Haifa, Israel; Wolfson Research Institute for Health and Wellbeing, Durham University, Durham, United Kingdom

**Author notes:** **E-Mail addresses:** RS; MS; GL; RDE; KB; PM; GRB. **Corresponding author:** Reinhard Stöger.

**Keywords:** Childhood, migrants, epigenetic age, RCP values, epigenetic stability, DNA methylation, accelerated ageing, Bangladesh, UK

## Abstract

**Background:** Migration from one environment to another often causes marked changes in developmental conditions. Here we compare epigenetic ageing and stability of the epigenetic maintenance system among British-Bangladeshi women who grew up in Bangladesh (adult migrants), where there are higher pathogen loads and poorer health care, to second-generation Bangladeshis who grew up in the UK. In our previous studies of these migrants, those who spent their childhoods in Bangladesh also had lower levels of reproductive hormones and a shorter reproductive lifespan compared to those who grew up in the UK, suggesting life history trade-offs during development. In the present study, we hypothesised that women who grew up in Bangladesh would have *i)* an older epigenetic/biological age compared to the women with a childhood in the UK and *ii)* that differences in the pace of epigenetic ageing might also be reflected by altered stability of DNA methylation marks.

**Results:** Illumina EPIC array methylation data from buccal tissue was used to establish epigenetic age estimates from 15 adult migrants and 11 second-generation migrants, aged 18-35 years. Using residuals from linear regression of DNA methylation-based biological age (DNAm age) on the chronological age, the results showed significant differences (p=0.016) in epigenetic age estimates: women whose childhood was in Bangladesh are on average 6.02 (± 2.34) years older, than those who grew up in London. We further investigated the efficiency of the epigenetic maintenance system which purportedly is reflected by epigenetic clocks. Methylation states of CpGs at the *LHCGR/LHR* locus, which contributes to Horvath’s multi tissue epigenetic clock were evaluated. Based on the Ratio of Concordance Preference (RCP) approach that uses double-stranded methylation data, we find that maintenance of epigenetic information is more stable in women who grew up in Bangladesh.

**Conclusions:** The work supports earlier findings that adverse childhood environments lead to phenotypic life history trade-offs. The data indicate that childhood environments can induce subtle changes to the epigenetic maintenance system that are detectable long after exposure occurred. The implication of such a finding warrants further investigation as it implies that a less flexible epigenetic memory system established early in life could reduce the capacity to respond to different environmental conditions in adult life.

## Background

Reproductive lifespans vary among individuals. Genetic variants associated with these complex traits, which include timing of puberty, age at first birth and age at menopause are closely related to fitness and undergo purifying selection [1,2]. The genetic architecture of reproductive ageing has been investigated largely in women of European ancestry. However, a limited number of studies in other populations suggests shared genetic underpinnings of these reproductive phenotypes, albeit with noticeable variations in effect allele frequencies and effect estimates in women of different ethnic groups [3–5]. Environmental exposures likely contribute to variations in heritability estimates and the phenotypic heterogeneity detected within and across different ethnic populations [6,7].

Our earlier work identified strong correlations between childhood environmental conditions and adult reproductive function [8–10]. In particular, Bangladeshi women who migrated as young adults to London, have lower levels of reproductive steroids when compared to British-Bangladeshi women who moved to the UK prior to the age of eight and women who were born in London to first-generation Bangladeshi immigrants [7,9–11]. An upbringing in Bangladesh is generally associated with a shortened reproductive lifespan, while its duration is longer for women with Bangladeshi ancestry, whose childhoods were spent in London [9]. Timing of reproductive functions across the life course correlates with the rate of ageing in other body systems [12].

Geographically and culturally the British-Bangladeshis women in these studies have a comparable background. They are all ethnic Bengalis and originally stem from a relatively affluent middle-class population in the northeast of Bangladesh and now live in East London. A possible environmental factor that distinguishes between the two childhood locations is the exposure to higher and recurrent infectious disease loads in Bangladesh [13–15]. Indeed, by mimicking early-life immune challenges in a mouse model, we replicated some of the distinct reproductive phenotypes characteristic of women with a childhood in Bangladesh, in including delayed onset of puberty lower ovarian reserve [16].

At the cellular level, environmental factors influence the chromatin state of the genome [17]. Stored as epigenetic information, cells have the capacity to retain some memory of past developmental and environmental conditions [18]. Methylation of genomic DNA is part of the epigenetic information storage system in mammalian cells where it is primarily confined to cytosines of CpG dinucleotides [19]. Methylation levels of discrete CpG sites have been used to develop remarkably accurate estimators of age. Such ‘epigenetic clocks’ link developmental and maintenance processes to biological ageing [reviewed in [20]]. Pace of ageing can vary and result in a mismatch between chronological and biological age of an individual [21].

Here, we explore the possible association between chronological age, biological ageing and an epigenetic maintenance system in Bangladeshi women of prime reproductive age (18-35 years old). The women of this study live within the same ethnic community in London but can be divided into two groups: those with a childhood in the UK, and those with a childhood in Sylhet, a city in the northeast region of Bangladesh. Using buccal cell DNA from these London-based Bangladeshi women, we recently identified genome-wide, altered DNA methylation levels between the two groups [16]. Since these DNA methylation measurements were generated on the MethylationEpic array platform, we re-examined the data using ‘Horvath’s clock’, a multi-tissue age-estimator with a robust relationship between chronological age and DNA methylation-based (DNAm) age [22].

## Results and Discussion

### Accelerated DNAm Age measured with Horvath’s epigenetic clock

We find that the correlation between chronological age and DNAm age does not differ significantly between women who grew up the UK (‘UK’ group; n=11) and women who grew up in Bangladesh (‘Bangladesh’ group; n=15). That is, chronological age affects DNAm age in a similar way in both groups (Additional file 1). However, regression analysis showed that the y-intercepts of the UK and Bangladeshi groups differ significantly (p=0.0083) / Additional file 1). This suggested that a childhood in Bangladesh correlates with DNAm Age predictions that differ noticeable when compared with epigenetic age estimates for women of the UK group.

The tick rate of epigenetic clocks is increased by many different environmental factors, including psychological traumas, smoking, asthma, alcohol, infections and hormonal changes following menopause [23–26]. Such acceleration of epigenetic age is best measured by residuals obtained from regressing DNAm age on chronological age [22]. Indeed, the pace of epigenetic ageing is accelerated in women with a childhood in Bangladesh and overall differs significantly from the UK group (p=0.016) (Figure 1). This altered pace of biological ageing is consistent with our previous observations that women who grow up in Bangladesh have a shorter reproductive lifespan and chronically lower levels of reproductive hormones [9,13,15]; reviewed in [7]. Although our finding of accelerated epigenetic ageing rests on a small number of sampled individuals, it highlights the limited utility of epigenetic clocks as a tool to determine the age – and consequently eligibility considerations – of asylum-seekers [27].

**Fig. 1.**
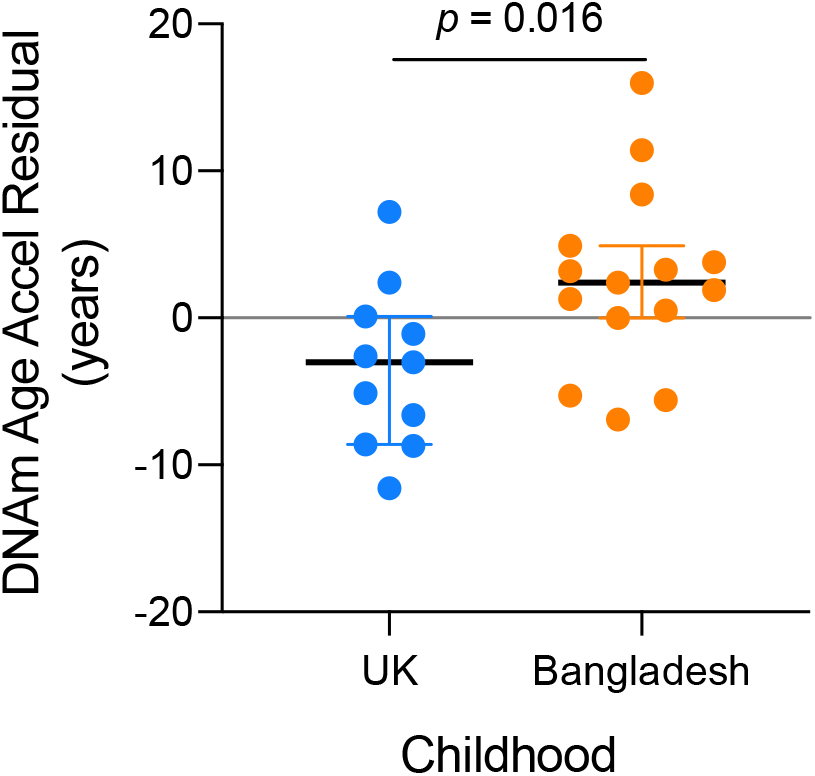
Differences in pace of epigenetic ageing. Plot of DNAm Age Accel Residuals, with each data point representing an individual. The colour indicates the corresponding dataset: blue = childhood in UK, orange = childhood in Bangladesh. The median is indicated by a horizontal line with upper and lower hinges representing the 25th and 75th percentiles. A positive or negative value indicates that the estimated epigenetic/biological age of the sample is higher or lower, respectively, than expected based on chronological age.

### Epigenetic stability of a clock locus

The tick rate of Horvath’s epigenetic clock is thought to reflect the rate at which work is done to maintain epigenetic stability [20,22]. It is possible to infer epigenetic stability by analysing double-stranded DNA methylation data with a new metric, Ratio of Concordance Preference (RCP) [28]. We used the RCP metric to estimate epigenetic stability at the *Luteinizing Hormone/Choriogonadotropin Receptor* (*LHCGR/LHR*) gene, which plays an important role in reproductive function. The *LHCGR* locus contains a CpG site, which contributes to Horvath’s DNAm Age clock [22].

We find that RCP estimates are generally higher for the ‘Bangladeshi’ group of women (Figure 2). Higher RCP estimates indicate higher levels of epigenetic stability [28]. That is, the methylation states of CpGs at *LHCGR* locus are more often identical on the two strands of individual DNA molecules of ‘Bangladeshi’ individuals when compared to ‘UK’ individuals. We note that the RCP estimates are based on a relatively small number of data points (Additional file 1), yet they are sufficient to indicate subtle differences in the workings of the epigenetic maintenance system between two groups of women who appear to age at different rates.

**Fig. 2.**
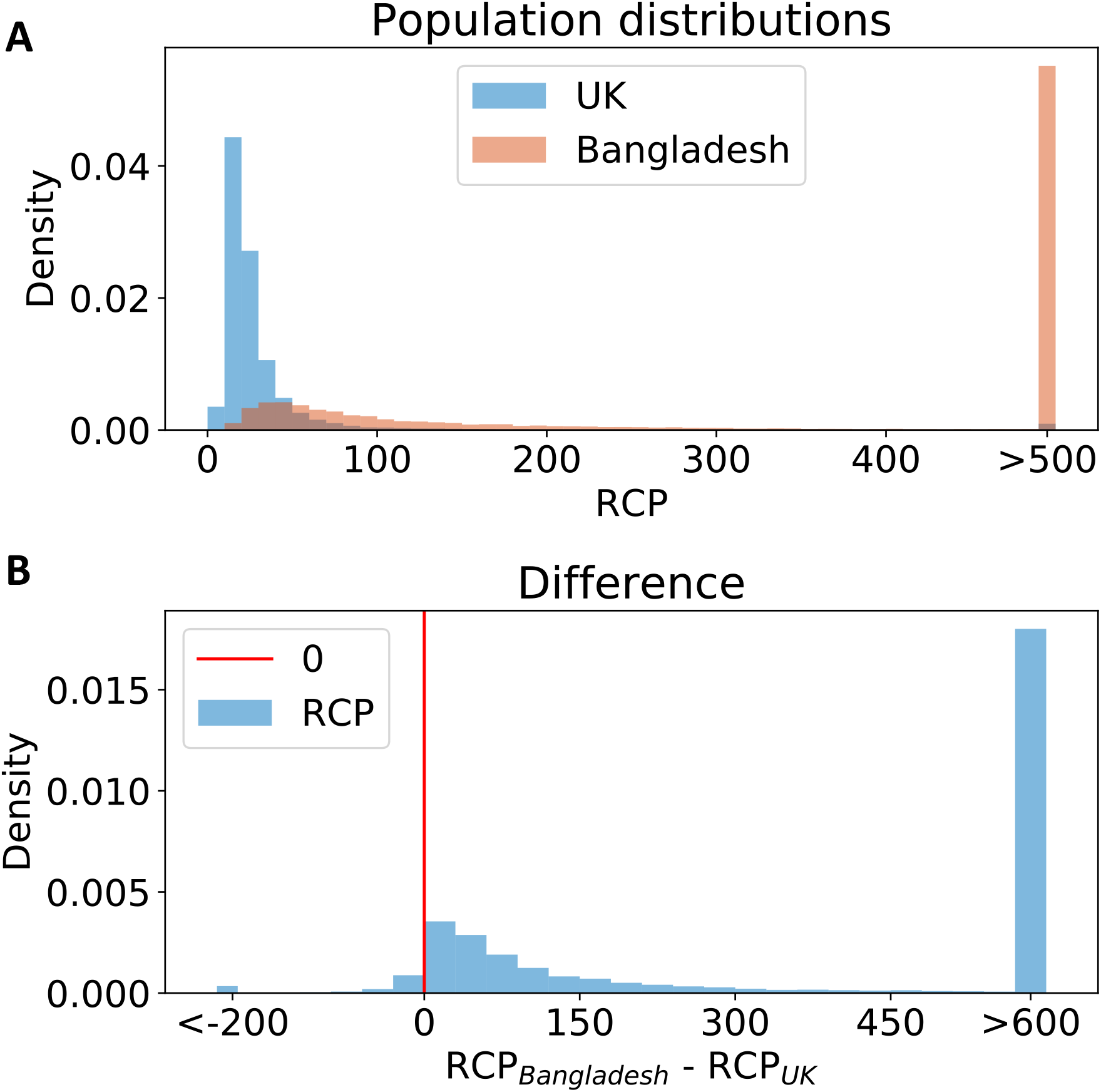
Inferences of DNA methylation stability differ between UK and Bangladeshi samples at the epigenetic-clock associated *LHCGR/LHR* locus. **A)** Ratio of Concordance Preference (RCP) is a metric that infers stability/flexibility of methylation states at matching CpG sites (CpG dyads) on the parent and daughter strand of individual DNA molecules, without assuming any specific enzymatic mechanisms of DNA methylation. Flexibility, indicated by RCP values near 1, indicates that the methylation system has no preference for either concordance or discordance of the methylation state at a CpG dyad and follows the random model. High RCP values – with the extreme approaching infinity – indicate high stability, where epigenetic maintenance systems have complete preference for concordant methylation states of CpG dyads (they are either methylated or unmethylated); none, or very few CpG dyads are hemi-methylated. Shown are the RCP distributions taken from bootstrap samples, weighing each individual evenly within each group (UK = blue, Bangladesh = orange). The sampled population of double-stranded DNA molecules – and the corresponding methylation states of CpG dyads – revealed a clear preference for a more stable epigenetic maintenance system in operation at the *LHR* locus in women with a childhood in Bangladesh, when compared to inferred RCP values for the samples from women with a childhood in London. **B)** Testing if the bootstrap samples of RCP differences (Bangladesh vs UK childhood) are significantly differnt. The red line is set at 0. The p-value is derived from this as the proportion of samples to the left of 0 (a one-tailed test to examine whether Bangladesh RCPs are significantly greater than UK RCP values). Two-tailed p-value is the double of that amount. p= 0.026 (one tailed); p= 0.052 (two tailed)

## Conclusions

The results of our study support a large body of work demonstrating phenotypic plasticity in response to environments encountered during early life. A childhood in Bangladesh measurably accelerates epigenetic/biological ageing in women, when compared to women of same chronological age (18-35 yrs) and ethnicity, who were born and brought up in London, UK. The multi tissue epigenetic clock is thought to register the workings of developmental and epigenetic maintenance systems linking these processes with the life course [20,22]. Our study is one of the first to test if differences in function of the epigenetic maintenance system can be linked with epigenetic age estimators. The findings indicate that subtle differences in the stability of epigenetic states are indeed associated with biological ageing and opens a new line of investigation.

## Methods

### DNA methylation data and establishment of DNAm Age

Genome-wide cytosine methylation levels were established using the Illumina HumanMethylationEPIC BeadChip Array following isolation of genomic DNA from buccal cells DNA (DNeasy Blood & Tissue Kit (Qiagen). Multidimensional scaling (MDS) plots indicated that no significant batch effects were skewing the MethylationEPIC BeadChip data sets. The data were processed with the Bioconductor/minfi package. CpG probes associated with known SNPs were removed, as were those with a detection probability of <0.01. Probes on both X and Y chromosomes were retained. Methylation beta values (0-1) were normalized by SWAN. The methylation data set (GSE133355 study) is accessible on the Gene Expression Omnibus (GEO) data platform at: https://www.ncbi.nlm.nih.gov/geo/query/acc.cgi?acc=GSE133355

### Determination of DNAm Age and age acceleration

A file with the beta values obtained from the Illumina HumanMethylationEPIC BeadChip Array work (see above) was used to establish the epigenetic age (DNAm age and AgeAccelerationResiduals) with Horvath’s method [22]. The underlying algorithms are available through the online DNA methylation calculator (http://dnamage.genetics.ucla.edu/).

### Generation of bisulfite hairpin data / methylation states of CpG dyads at the *LHCGR/LHR* locus

We have previously described in detail the concept and procedure of generating authenticated, non-redundant double-stranded DNA methylation data [19,29,30]. In brief, genomic sequence information surrounding the *LHR* clock-CpG site [one of the 353 CpG sites contributing to Horvath’s clock [22]; Illumina cluster ID cg12351433 / chr2:48982957-48982957 / UCSC Genome Browser (GRCh37/hg19)] was used to identify suitable restriction recognition sites to generate 3’, or 5’-overhangs, respectively, for the ligation of UMI-barcoded hairpin linkers. Specifically, restriction enzymes StyI or BstXI (New England Biolabs) were used. Combinations of the following primers were used to amplify hairpin-linked, bisulfite converted DNA:

bsLHR-R1 5’-RCAAATCAAAACAAAACAAACTC-3’;
bsLHR-R2 5’-CACTAAACACTATCRCAAATCAAAAC-3’;
bsLHR-F1 5’-TAGTAGGAAGGAGGTTATTGG-3’;
bsLHR-F2 5’-GTAGGTTAAGGTAGAGTAGATTTAG-3’;
bsLHR-F3 5’-GAATTGGGTTTTTGCGGTTTGTTAG-3’.

Further information of the hairpin-concept and of the barcoded and batch-stamped hairpin linkers (Eurofins Genomics) are provided in Additional file 1.

#### Processing of the sequencing data

Fold is a web application for the analysis of the output of hairpin-bisulphite sequencing data. Specifically, the programme reconstructs, visualises, and generates statistics on the double-stranded CpG methylation patterns of the original cohort of DNA molecules. This is achieved by first ‘realigning’ the top and bottom strand of the molecule about the hairpin, in which the programme attempts to manage ‘PCR slippage’, and other sequencing errors. Then algorithm then identifies and categorises CpG dyads, which is possible due to the previous bisulphite conversion of unmethylated cytosine to uracil (and so recognised as tyrosine when sequenced). For example, fully methylated dyads are those regions in where the reconstructed top strand is C-G and the bottom is G-C. Similarly, fully unmethylated dyads are those where the top is T-G and the bottom is G-T. In addition, the programme calculates a metric: ‘Ratio of Concordance Principle’ which quantifies the concordance of methylation between the top and bottom strands of the DNA molecule (0=complete discordance, 1=random concordance, inf=complete concordance). This metric represents the preference of the summation of epigenetic mechanisms of the cell to either maintain or obscure methylation patterns of the DNA in the cells at the time the sample was taken. The functions of Fold was written in R and the web application is written in PHP. The live web application can be found at http://www.gregoryleeman.com/fold, and the repository can be found at https://github.com/gregoryleeman/fold.

### Analysis and comparison of RCPs at the *LHCGR/LHR* locus

RCP values are based on double-stranded DNA methylation data derived from sequences of individual hairpin bisulfite PCRs products. RCP analyses were done following the procedures described in [28] with the small addition of bootstrapping individuals within each population. The additional step in the procedure helps to address the possibility of uneven sampling from a larger population. The analysis procedures in brief are described below. Each population RCP distribution was drawn through hierarchical bootstrap sampling. For each of 20,000 bootstrap samples, individuals in each population were sampled with replacement, and double stranded DNA sequences of each of the sampled individuals were in turn sampled with replacement. Dyad counts were then normalised such that each individual had the same number of dyads. The normalised dyad counts were then summed, corrected for failed bisulfite conversions (rate of 0.0039, measured empirically) and inappropriate conversions (rate of 0.017, estimated as described in [31] [Genereux et al., 2008]), and used to compute the RCP value. A bootstrap sample of the RCP difference was computed by taking the difference of the RCP values sampled for the two populations.

For one-tailed comparison tests, with which we examine directional differences, we determined the p-value as the proportion of bootstrap-difference samples to the left of 0. For two-tailed tests, with which we can detect differences in any direction, we determined the p-value as twice the smaller proportion of the bootstrap difference samples on either side of 0.

## Availability of data and materials

The datasets generated and/or analysed during the current study are available in the Gene Expression Omnibus (GEO) data platform https://www.ncbi.nlm.nih.gov/geo/query/acc.cgi?acc=GSE133355

All data generated or analysed during this study are included in this published article and its supplementary information file.

## Authors’ contributions

RS: Conceptualisation, Funding acquisition, Experimental work, Analysis, Resources, Supervision, Data curation, Project administration, Writing – original draft, Writing – review & editing.

MC: Analysis, review & editing

GL: Analysis, Coding

RDE: Analysis, Data curation

KB: Experimental work, Resources

PM: Funding acquisition, Project administration, Writing – review & editing.

GRB: Conceptualisation, Funding acquisition, Project administration, Writing – review & editing.

## Acknowledgements

We thank Kamila Derecka for technical support, Ian C. W. Hardy for statistical advice and Steve Horvath for information on DNAm age analysis.

## Additional file 1: Chronological age vs DNAm Age

### Estimates of DNA methylation age (DNAm Age)

The online calculator (https://dnamage.genetics.ucla.edu/) was used to generate estimates of DNAm Age, AgeAccelerationDiff (= DNAmAge-Age), and AgeAccelerationResidual (= the recommended age acceleration measure based on Horvath’s linear regression model [Horvath S (2013) DNA methylation age of human tissues and cell types. *Genome Biol 14*(10):R115 PMID: 24138928].

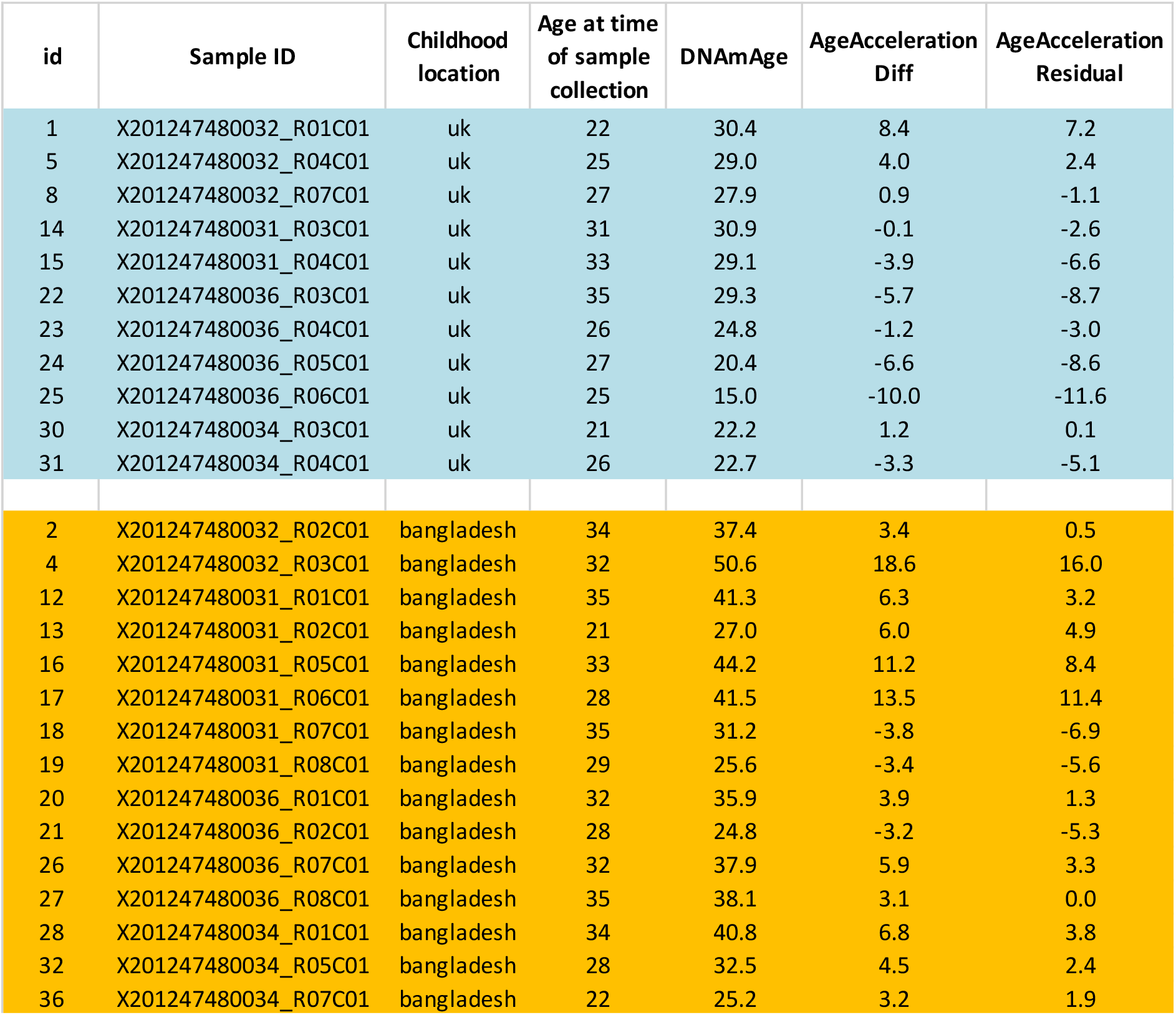

| id | Sample ID | Childhood location | Age at time of sample collection | DNAmAge | AgeAccelerationDiff | AgeAccelerationResidual |
| --- | --- | --- | --- | --- | --- | --- |
| 1 | X201247480032_R01C01 | uk | 22 | 30.4 | 8.4 | 7.2 |
| 5 | X201247480032_R04C01 | uk | 25 | 29.0 | 4.0 | 2.4 |
| 8 | X201247480032_R07C01 | uk | 27 | 27.9 | 0.9 | -1.1 |
| 14 | X201247480031_R03C01 | uk | 31 | 30.9 | -0.1 | -2.6 |
| 15 | X201247480031_R04C01 | uk | 33 | 29.1 | -3.9 | -6.6 |
| 22 | X201247480036_R03C01 | uk | 35 | 29.3 | -5.7 | -8.7 |
| 23 | X201247480036_R04C01 | uk | 26 | 24.8 | -1.2 | -3.0 |
| 24 | X201247480036_R05C01 | uk | 27 | 20.4 | -6.6 | -8.6 |
| 25 | X201247480036_R06C01 | uk | 25 | 15.0 | -10.0 | -11.6 |
| 30 | X201247480034_R03C01 | uk | 21 | 22.2 | 1.2 | 0.1 |
| 31 | X201247480034_R04C01 | uk | 26 | 22.7 | -3.3 | -5.1 |
| 2 | X201247480032_R02C01 | bangladesh | 34 | 37.4 | 3.4 | 0.5 |
| 4 | X201247480032_R03C01 | bangladesh | 32 | 50.6 | 18.6 | 16.0 |
| 12 | X201247480031_R01C01 | bangladesh | 35 | 41.3 | 6.3 | 3.2 |
| 13 | X201247480031_R02C01 | bangladesh | 21 | 27.0 | 6.0 | 4.9 |
| 16 | X201247480031_R05C01 | bangladesh | 33 | 44.2 | 11.2 | 8.4 |
| 17 | X201247480031_R06C01 | bangladesh | 28 | 41.5 | 13.5 | 11.4 |
| 18 | X201247480031_R07C01 | bangladesh | 35 | 31.2 | -3.8 | -6.9 |
| 19 | X201247480031_R08C01 | bangladesh | 29 | 25.6 | -3.4 | -5.6 |
| 20 | X201247480036_R01C01 | bangladesh | 32 | 35.9 | 3.9 | 1.3 |
| 21 | X201247480036_R02C01 | bangladesh | 28 | 24.8 | -3.2 | -5.3 |
| 26 | X201247480036_R07C01 | bangladesh | 32 | 37.9 | 5.9 | 3.3 |
| 27 | X201247480036_R08C01 | bangladesh | 35 | 38.1 | 3.1 | 0.0 |
| 28 | X201247480034_R01C01 | bangladesh | 34 | 40.8 | 6.8 | 3.8 |
| 32 | X201247480034_R05C01 | bangladesh | 28 | 32.5 | 4.5 | 2.4 |
| 36 | X201247480034_R07C01 | bangladesh | 22 | 25.2 | 3.2 | 1.9 |

### Prism GraphPad 8 analysis / Chronological age vs DNAm Age

The slopes of the regression lines between the ‘UK’ group (n=11) and the ‘Bangladesh’ group (n=15) are not significantly different (F = 1.136; DFn = 1, DFd = 22 p=0.29). Therefore, a single slope for data of the entire cohort can be established: the pooled slope equals 0.8128 (Additional Fig 1a). However, the intercepts are significantly different (F = 8.341. DFn = 1, DFd = 23, p=0.0083)

**Additional Fig 1a:**
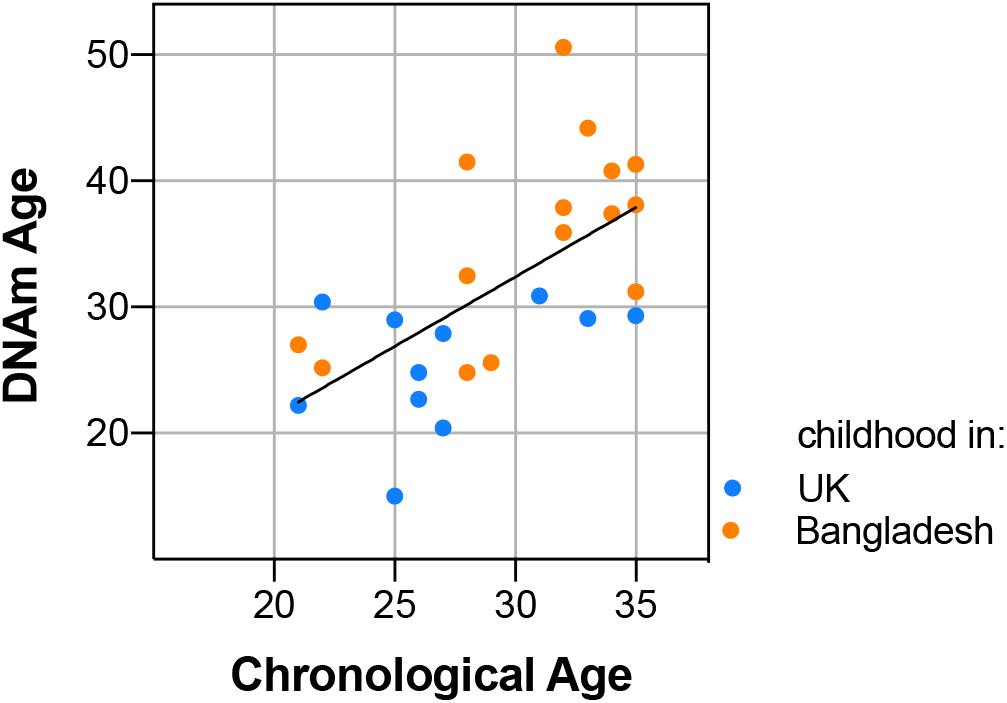
Plot of predicted methylation age (Horvath clock) against chronological age. The scatter plot shows DNA methylation age vs. chronological age vs. and the line in which DNA methylation age was regressed on chronological age using both, the ‘UK’ and the ‘Bangladesh’ data sets. ‘UK’ = Bangladeshi women who grew up in London, UK (blue); ‘Bangladesh’ = Bangladeshi women who grew up in Sylhet, Bangladesh. Each data point represents an individual, with the colour indicating the corresponding dataset.

### Genstat analysis / Chronological age vs DNAm Age

Genstat analysis of covariance yielded similar results to those obtained by Prism GraphPad 8 analysis, in that Bangladeshi women with a childhood in the UK and Bangladeshi women with a childhood in Bangladesh are affected by chronological age the same way – the two slopes are the same based on a minimal adequate mode (Additional Fig 1b). Parameters needed to reconstruct the lines are indicated in yellow:

Estimates of parameters
| Parameter | estimate | s.e. | t(23) | t pr. |
| --- | --- | --- | --- | --- |
| Constant | 10.78 | 8.35 | 1.29 | 0.209 |
| Country UK | -7.19 | 2.49 | -2.89 | 0.008 |
| chronological_age | 0.813 | 0.269 | 3.02 | 0.00 |

Parameters for factors are differences compared with the reference level:
| Factor | Reference level |
| --- | --- |
| Country | Bangladesh |

Accumulated analysis of variance
| Change | d.f. | s.s. | m.s. | v.r. | F pr. |
| --- | --- | --- | --- | --- | --- |
| + chronological_age | 1 | 660.09 | 660.09 | 19.57 | <.001 |
| + Country | 1 | 282.95 | 282.95 | 8.39 | 0.008 |
| + chronological_age.Country | 1 | 38.32 | 38.32 | 1.14 | 0.298 |
| Residual | 22 | 741.94 | 33.72 |  |  |
| - chronological_age.Country | -1 | -38.32 | 38.32 | 1.14 | 0.298 interaction not significant |
| - Country | -1 | -282.95 | 282.95 | 8.39 | 0.008 country significant |
| + Country | 1 | 282.95 | 282.95 | 8.39 | 0.008 country put back into model |
| - chronological_age | -1 | -309.57 | 309.57 | 9.18 | 0.006 age significant |
| + chronological_age | 1 | 309.57 | 309.57 | 9.18 | 0.006 age put back into model, so that the final model can be plotted. |
| Total | 25 | 1723.29 |  | 68.93 |  |

**Additional Fig 1b:**
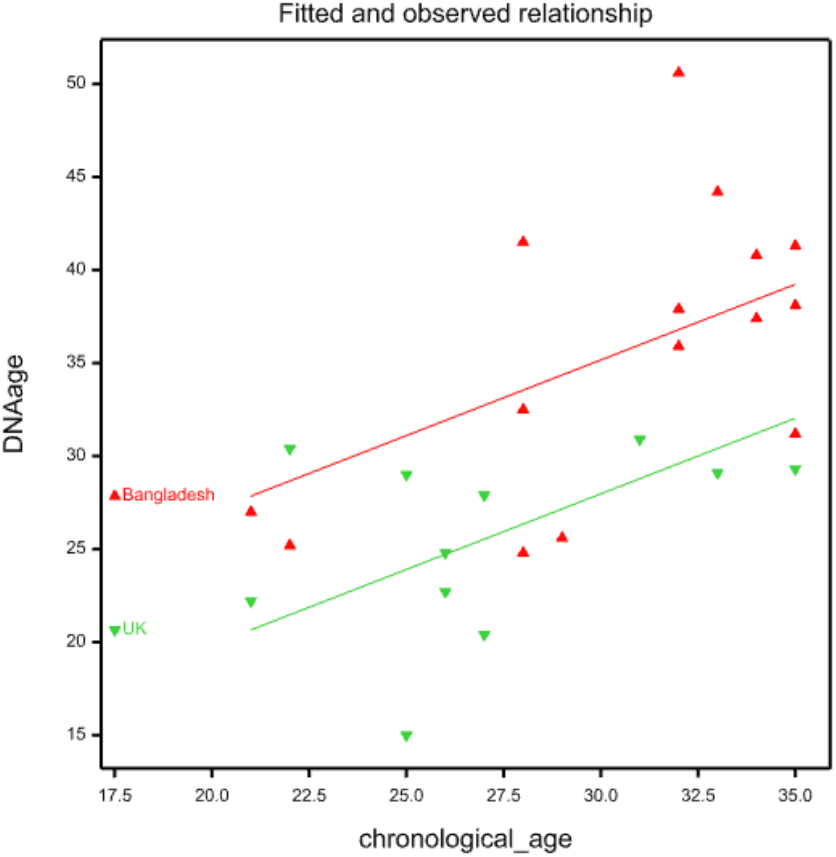
The minimal adequate model (Genstat analysis):

### DNA sequence surrounding the *LHR*-clock CpG site analysed by hairpin bisulfite PCR

The CpG site [clock CpG] at the *Luteinizing Hormone/Choriogonadotropin Receptor* (*LHCGR/LHR*) locus, which contributes to ‘Horvath’s clock’ (Horvath, 2013) was PCR-amplified following hairpin linker-ligation and sodium bisulfite conversion. The hairpin PCR products also captured methylation information of flanking CpGs (highlighted in red): four CpGs with the StyI-hairpin linker approach and eight CpGs with the BstXI-hairpin linker approach, respectively:

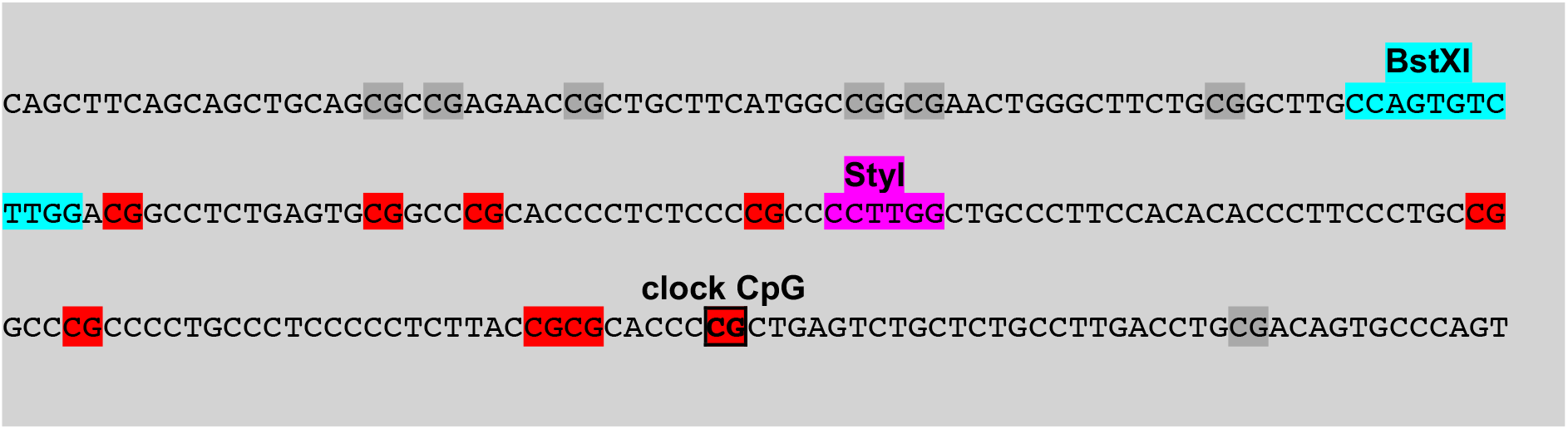

### BstXI- and StyI-hairpin linkers used in this study

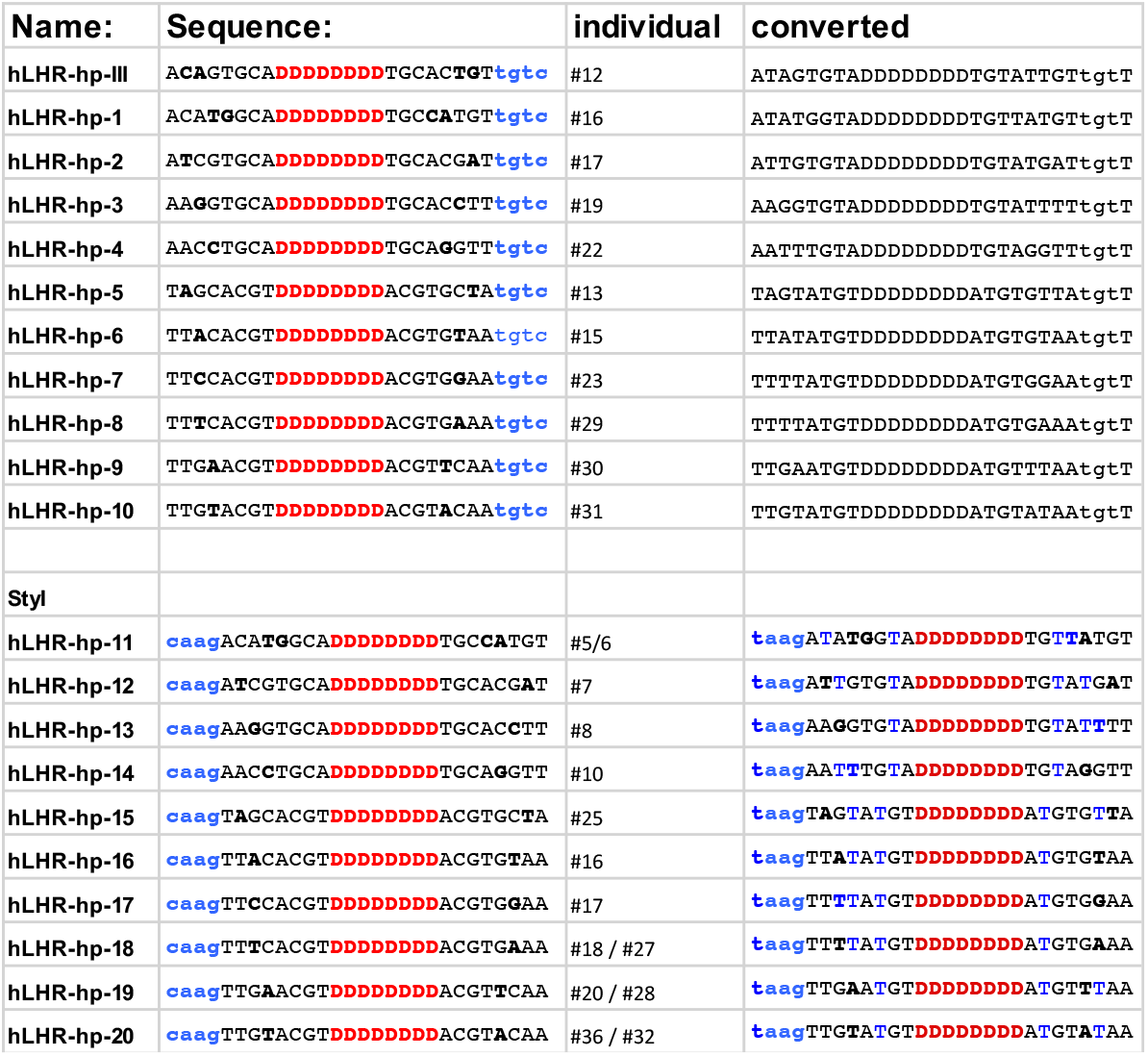

### Double-stranded methylation data of individual DNA molecules derived from buccal cells

The data shown on page 5 are processed, in that the methylation status of matching CpG sites of the top- and bottom strands (CpG dyads) are indicated as methylated (=1), or unmethylated (=0). Information of individual, double-stranded DNA molecules is displayed as follows:

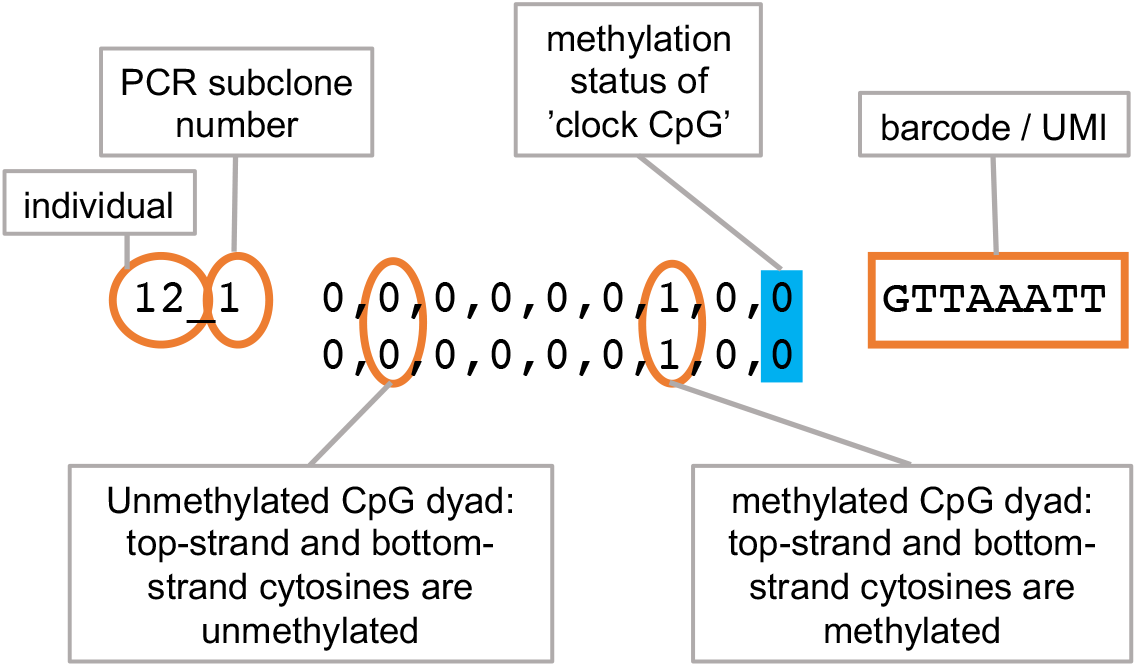

### Hairpin methylation data / Bangladesh-childhood

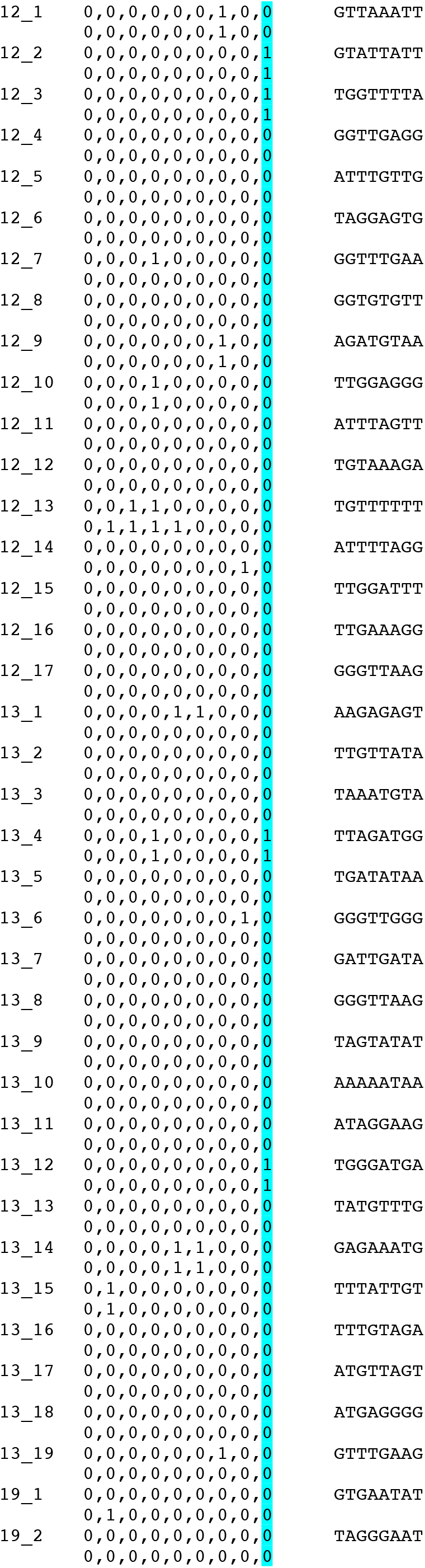
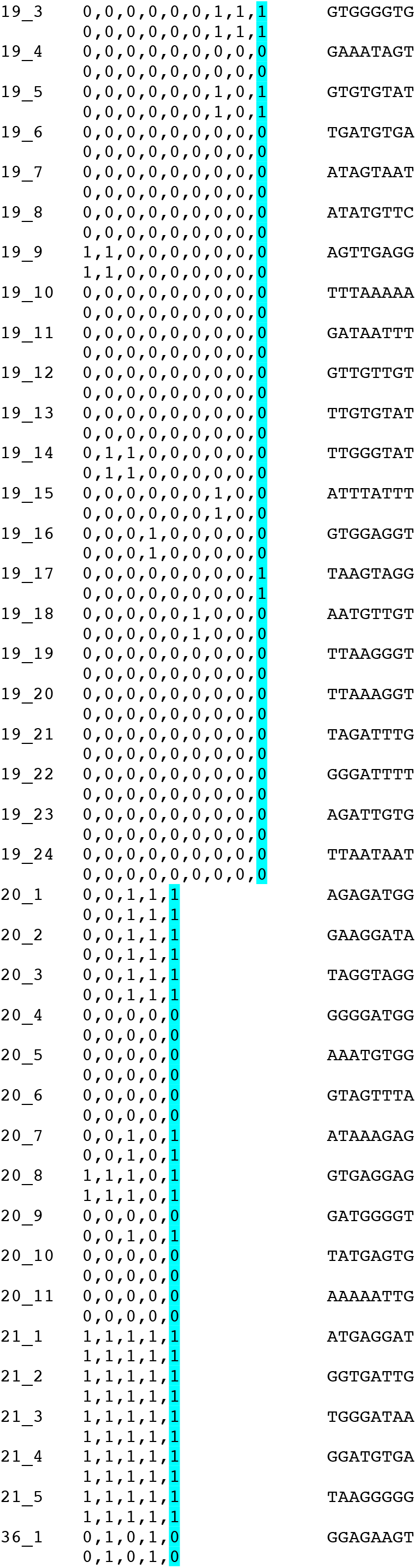
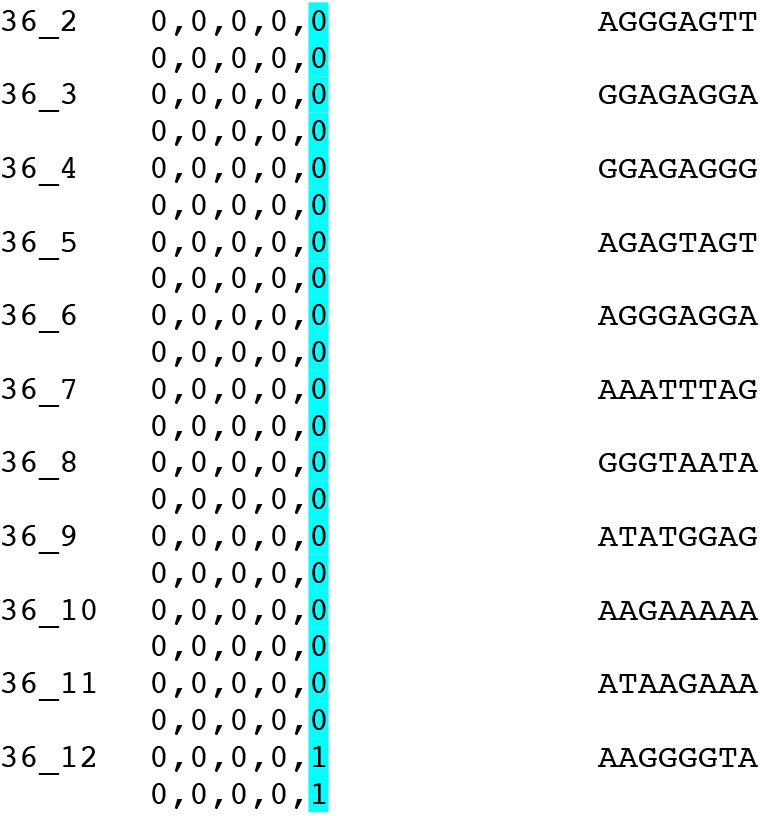

### Hairpin methylation data / Bangladesh-childhood

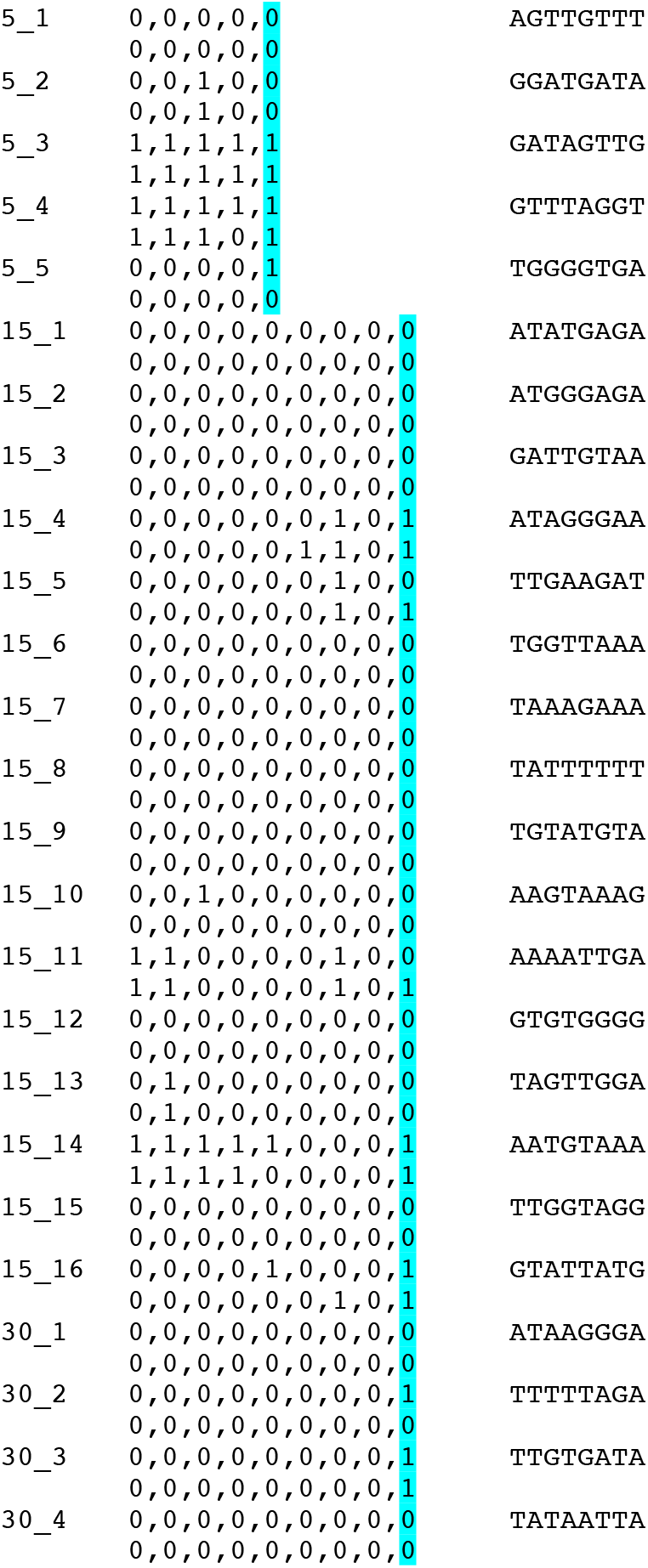
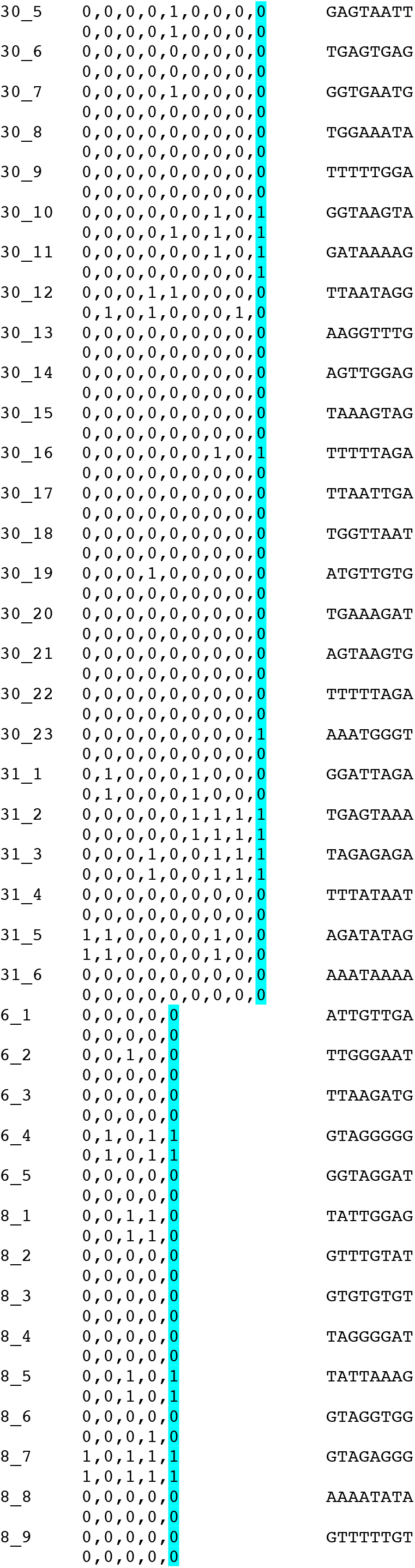
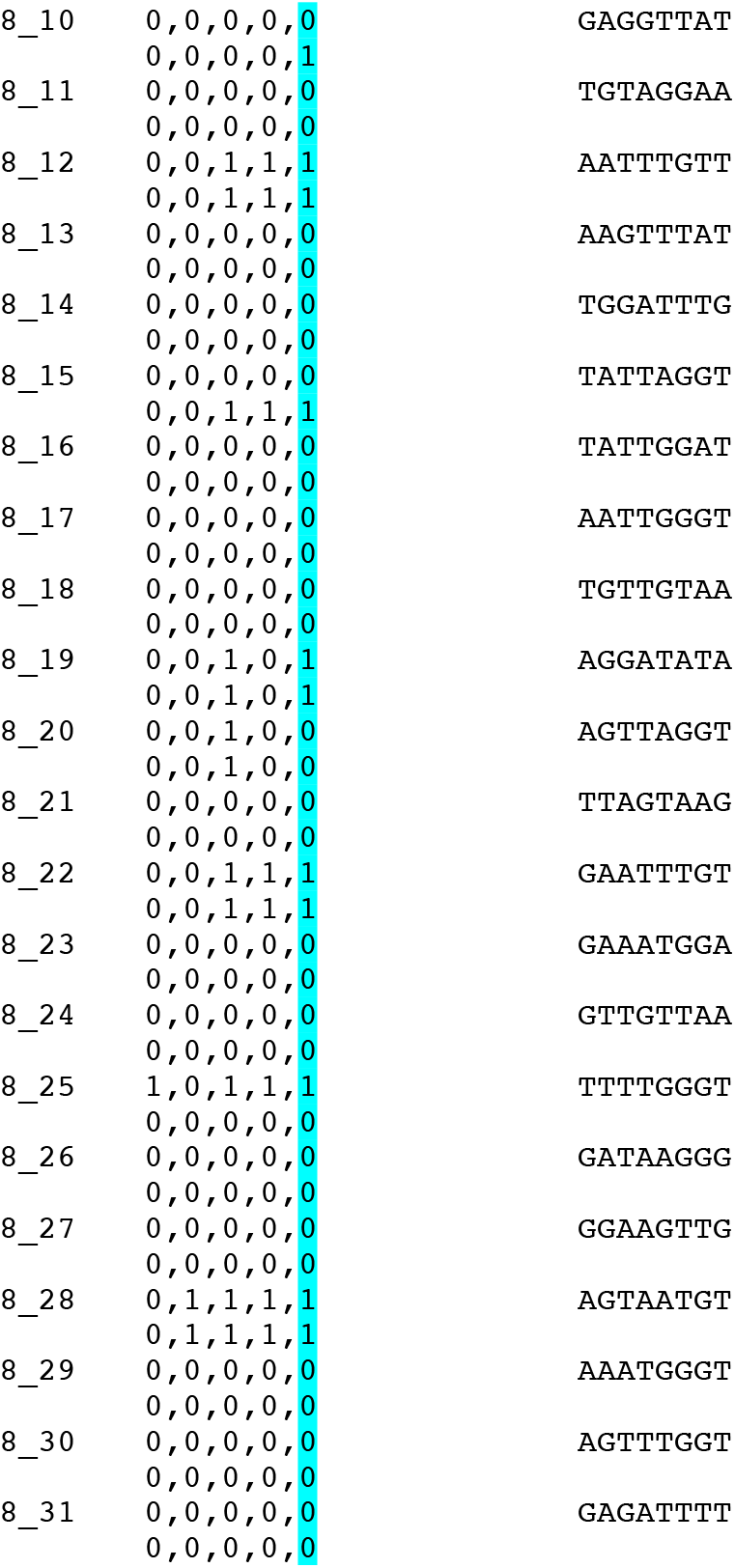

## References

1. Day FR, Ruth KS, Thompson DJ, Lunetta KL, Pervjakova N, Chasman DI, et al. Large-scale genomic analyses link reproductive aging to hypothalamic signaling, breast cancer susceptibility and BRCA1-mediated DNA repair. Nat Genet. Nature Publishing Group; 2015;47:1294–303.

2. Zeng J, De Vlaming R, Wu Y, Robinson MR, Lloyd-Jones LR, Yengo L, et al. Signatures of negative selection in the genetic architecture of human complex traits. Nat Genet. Nature Publishing Group; 2018;50:746–53.

3. Fernández-Rhodes L, Malinowski JR, Wang Y, Tao R, Pankratz N, Jeff JM, et al. The genetic underpinnings of variation in ages at menarche and natural menopause among women from the multi-ethnic Population Architecture using Genomics and Epidemiology (PAGE) Study: A trans-ethnic meta-analysis. Thameem F, editor. PLoS One. Public Library of Science; 2018;13:e0200486.

4. Horikoshi M, Day FR, Akiyama M, Hirata M, Kamatani Y, Matsuda K, et al. Elucidating the genetic architecture of reproductive ageing in the Japanese population. Nat Commun. Nature Publishing Group; 2018;9:1–9.

5. Carty CL, Spencer KL, Setiawan VW, Fernandez-Rhodes L, Malinowski J, Buyske S, et al. Replication of genetic loci for ages at menarche and menopause in the multi-ethnic Population Architecture using Genomics and Epidemiology (PAGE) study. Hum Reprod. Oxford University Press; 2013;28:1695–706.

6. Parent AS, Teilmann G, Juul A, Skakkebaek NE, Toppari J, Bourguignon JP. The Timing of Normal Puberty and the Age Limits of Sexual Precocity: Variations around the World, Secular Trends, and Changes after Migration. Endocr. Rev. Endocr Rev; 2003. p. 668–93.

7. Bar-Sadeh B, Rudnizky S, Pnueli L, Bentley GR, Stöger R, Kaplan A, et al. Unravelling the role of epigenetics in reproductive adaptations to early-life environment. Nat. Rev. Endocrinol. Nature Research; 2020. p. 519–33.

8. Magid KS, Uddin Ahamed F, Lawson DW, Chatterton RT, Bentley GR. Effects of adult migration on male salivary testosterone. Am J Hum Biol. 2006;18:262.

9. Begum K, Muttukrishna S, Sievert L, Sharmeen T, Murphy L, Chowdhury O, et al. Ethnicity or environment: Effects of migration on ovarian reserve among Bangladeshi women in the UK. Fertil Steril. 2015;under revi.

10. Nunez-De La Mora A, Bentley GR, Choudhury OA, Napolitano DA, Chatterton RT. The impact of developmental conditions on adult salivary estradiol levels: why this differs from progesterone? Am J Hum Biol. 2007/10/25. 2008;20:2–14.

11. Houghton LC, Cooper GD, Booth M, Chowdhury OA, Troisi R, Ziegler RG, et al. Childhood environment influences adrenarcheal timing among first-generation Bangladeshi migrant girls to the UK. Sear R, editor. PLoS One. 2014/10/14. 2014;9:e109200.

12. Perry JRB, Murray A, Day FR, Ong KK. Molecular insights into the aetiology of female reproductive ageing. Nat. Rev. Endocrinol. Nature Publishing Group; 2015. p. 725–34.

13. Núñez-de la Mora A, Chatterton RT, Choudhury OA, Napolitano DA, Bentley GR, Nunez-de la Mora A, et al. Childhood conditions influence adult progesterone levels. Fisk NM, editor. PLoS Med. 2007/05/17. 2007;4:e167.

14. Siddique AK, Baqui AH, Eusof A, Zaman K. 1988 floods in Bangladesh: pattern of illness and causes of death. J Diarrhoeal Dis Res. 1991;9:310–4.

15. Murphy L, Sievert L, Begum K, Sharmeen T, Puleo E, Chowdhury O, et al. Life course effects on age at menopause among Bangladeshi sedentees and migrants to the UK. Am J Hum Biol. 2012/11/24. 2013;25:83–93.

16. Bar-Sadeh B, Pnueli L, Begum K, Leeman G, Emes RD, Stöger R, et al. Early-life environment programs reproductive strategies through epigenetic regulation of SRD5A1 1. bioRxiv. Cold Spring Harbor Laboratory; 2020;2020.09.16.299560.

17. Zhu J, Adli M, Zou JY, Verstappen G, Coyne M, Zhang X, et al. Genome-wide Chromatin State Transitions Associated with Developmental and Environmental Cues. Cell. 2013;152:642–54.

18. Jaenisch R, Bird A. Epigenetic regulation of gene expression: how the genome integrates intrinsic and environmental signals. Nat Genet. 2003;33 Suppl:245–54.

19. Laird CD, Pleasant ND, Clark AD, Sneeden JL, Hassan KMA, Manley NC, et al. Hairpin-bisulfite PCR: Assessing epigenetic methylation patterns on complementary strands of individual DNA molecules. Proc Natl Acad Sci U S A. National Academy of Sciences; 2004;101:204–9.

20. Horvath S, Raj K. DNA methylation-based biomarkers and the epigenetic clock theory of ageing. Nat Rev Genet. Nature Publishing Group; 2018;19:371–84.

21. Khan SS, Singer BD, Vaughan DE. Molecular and physiological manifestations and measurement of aging in humans. Aging Cell. Blackwell Publishing Ltd; 2017;16:624–33.

22. Horvath S. DNA methylation age of human tissues and cell types. Genome Biol. 2013/10/22. BioMed Central Ltd; 2013;14:R115.

23. Dhingra R, Nwanaji-Enwerem JC, Samet M, Ward-Caviness CK. DNA Methylation Age—Environmental Influences, Health Impacts, and Its Role in Environmental Epidemiology. Curr. Environ. Heal. reports. Springer; 2018. p. 317–27.

24. Quach A, Levine ME, Tanaka T, Lu AT, Chen BH, Ferrucci L, et al. Epigenetic clock analysis of diet, exercise, education, and lifestyle factors. Aging (Albany NY). 2017;9:419–46.

25. Austin MK, Chen E, Ross KM, McEwen LM, Maclsaac JL, Kobor MS, et al. Early-life socioeconomic disadvantage, not current, predicts accelerated epigenetic aging of monocytes. Psychoneuroendocrinology. 2018;97:131–4.

26. Levine ME, Lu AT, Quach A, Chen BH, Assimes TL, Bandinelli S, et al. An epigenetic biomarker of aging for lifespan and healthspan. Aging (Albany NY). 2018;10:573–91.

27. Editorial. Molecular test of age highlights difficult questions. 2018;561.

28. Choi M, Genereux DP, Goodson J, Al-Azzawi H, Allain SQ, Simon N, et al. Epigenetic memory via concordant DNA methylation is inversely correlated to developmental potential of mammalian cells. Barsh GS, editor. PLOS Genet. 2017;13:e1007060.

29. Miner BE, Stöger RJ, Burden AF, Laird CD, Hansen RS. Molecular barcodes detect redundancy and contamination in hairpin-bisulfite PCR. Nucleic Acids Res. 2004;32.

30. Stöger R. Hairpin-Bisulfite PCR. Methods Mol Biol. NLM (Medline); 2021;2198:287–99.

31. Genereux DP, Johnson WC, Burden AF, Stöger R, Laird CD. Errors in the bisulfite conversion of DNA: modulating inappropriate- and failed-conversion frequencies. Nucleic Acids Res. 2008/11/06. Oxford University Press; 2008;36:e150.

